# Computer models predict differential dendritic vulnerability with ischemia and spreading depression

**DOI:** 10.1101/2025.09.22.677786

**Authors:** Adam J.H. Newton, William W. Lytton, Marcello DiStasio, Robert A. McDougal

## Abstract

Ischemia, whether abrupt or chronic, limits ATP production and disrupts ATP-dependent homeostatic mechanisms, leading to alterations in both intracellular and extracellular ion concentrations. Inadequate neuronal ATP triggers K^+^ release and increased extracellular K^+^ depolarizes neurons, leading to additional K^+^ release; this positive feedback phenomenon is known as spreading depolarization (SD). When the depolarizing effects are strong enough, the cells undergo depolarization blockade, known as spreading depression. Excess extracellular K^+^ increases energy demand from the Na^+^-K^+^ pump, producing a pathological confluence of increased demand with reduced delivery of energy. The resulting changes have profound effects at subcellular, cellular, and network scales of brain function. We hypothesized that consequences of ischemic or SD homeostatic failure would differ on the subcellular scale, with differences between disjunct dendritic regions of a hippocampal CA1 pyramidal neuron. To evaluate the interplay between morphology and ion concentrations, we used a mechanistic simulation incorporating neuronal morphology, pumps, exchangers, voltage-, and Ca^2+^-sensitive ion channels. In both cases, calcium accumulation was greatest in the basilar dendrites, suggesting these dendrites would show the greatest effects of excitotoxicity. By contrast, ischemia, but not SD showed that distal apical dendrites were exposed to greater intracellular chloride concentrations, which may lead to dendritic beading.

## 1 Introduction

Spreading depolarization (SD), seen in a wide range of neurological diseases, is a wave of neuronal depolarization associated with changes in extracellular ionic concentrations, particularly in K^+^ and Na^+^. SD propagates through neural tissue at ∼ 1−8 mm/min (Drenckhahn et al., 2012; Hartings et al., 2017; Dreier and Reiffurth, 2015). The changes due to SD are temporary, but restoring ionic concentrations requires energy. Therefore, a pathological synergy occurs: SD augments local ischemia, and ischemia prevents recovery of SD. Furthermore, transient ischemia will trigger SD and worsen ischemia. Spatiotemporal interactions between SD overuse and oxygen underdelivery explain some of the mottled or heterogeneous tissue damage seen in patients with a variety of disorders associated with SD. Notably, chronic migraineurs show anomalies on MRI suggesting episodic tissue damage(Hadjikhani et al., 2001). SD is also associated with traumatic brain injury, tumor, epilepsy, and ischemic stroke and can be associated with MRI anomalies in all of these disorders, sometimes far from the primary site of injury Dreier (2011).

Continuous oxygen delivery is essential to produce the ATP needed to maintain ion homeostasis and maintain tissue viability, providing energy for intracellular and neuron-to-neuron signaling that drive brain function(Hall et al., 2012). Most of the energy consumption of neural tissue goes to the Na^+^-K^+^ pump to drive these ions against their gradients: three [Na^+^]_i_ are exchanged for two [K^+^]_o_(Attwell and Laughlin, 2001). The time course of an energy deficit at the pump will determine the risk of cell death and the nature of cell damage (Dreier, 2011; Risher et al., 2010). Energy consumed by the Na^+^-K^+^ pump drives other exchangers, which depend on Na^+^ and K^+^ ion gradients, although some may also consume ATP directly.

Overall, ionic homeostasis maintains a variety of gradients responsible for sustaining various aspects of cell function, which may fail at different times and to different extents. Varying spatiotemporal failure patterns will produce distinct patterns of ion failure, with different subcellular and tissue pathology patterns. Na^+^ and K^+^ gradients are required for action potential generation and transmission. The Ca^2+^ gradient is required for synaptic transmission (Somjen, 2004), using a gradient actively maintained by two ATP pumps, plasma membrane Ca^2+^ ATPase (PMCA), and the sarco-endoplasmic reticulum Ca^2+^ ATPase (SERCA). These Ca^2+^ pumps have far less energy demand than the Na^+^-K^+^ pump as the Ca^2+^ gradient is smaller (Somjen, 2001);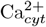 is also extruded from the neuron by Na^+^-Ca^2+^ exchange, utilizing the Na^+^ gradient. Cl^−^ symporters exploit the gradient created by the Na^+^ -K^+^ pump, without directly consuming any ATP, to move Cl^−^ in or out of the cell (Watanabe and Fukuda, 2015).

The ubiquity of SD in multiple pathologies indicates its relevance as a target for a variety of particularized clinical interventions, avoiding a threatened pattern and pace of damage expected from predicted pathophysiology(Gorji, 2001). Monitoring the frequency and duration of SDs following growth of a tumor, an ischemic or hemorrhagic stroke, or other expanding lesion can serve as a biomarker for the metabolic state of the tissue and suggest secondary therapy (Dreier et al., 2017).

Before causing cell death, ischemia and SD lead to cell damage, manifested histologically as dendritic beading, where water enters – partially through Cl^−^ symporters (NKCC1 and KCC2) (Steffensen et al., 2015; Hellas and Andrew, 2021) – causing varicosities and spine loss, or outright dendritic loss (Risher et al., 2010). Dendritic loss can occur late following beading Hoskison and Shuttleworth (2006); Hasbani et al. (1998) or may occur more immediately due to Ca^2+^ entry causing excitotoxicity. The microanatomical patterns of dendritic damage, loss, and of cell death depend on the details of the underlying causes.

We hypothesized that SD and ischemia, in various combinations of severity, will lead to predictable spatiotemporal patterns of damage which can predict the extent of prolonged and extended homeostatic failure and thus guide therapy. To test this, we used a model pyramidal cortical neuron with detailed morphology, intracellular Ca^2+^ dynamics (Migliore et al., 2018; Neymotin et al., 2014), glutamate activation (Hübel et al., 2017) and ion pumps (Wei et al., 2014; Le Masson et al., 2014). We also included a single simplified astrocyte cell model, represented as a line of compartments parallel to the neuron model. The astrocyte model also consumed energy to clear glutamate (Hübel et al., 2017) and K^+^ (Somjen et al., 2009; Kager et al., 2002, 2000).

The model parameters were tuned to produce physiological spike rates and maintain homeostasis under simulated current clamp. Subsequently, we simulated three conditions: (1) elevated [K^+^]_o_, causing SD alone as would be seen in migraine with aura; (2) hypoxia/ischemia at baseline [K^+^]_o_, as in ischemic tissue that may generate secondary SD; and (3) ischemia with elevated [K^+^]_o_, as in hypoxic spreading depolarization (HSD). HSD exhibited similar dynamics as SD over short timescales and showed behavior that converged with ischemia over longer timescales.

We then focused on two contrasting cases, (i) SD and (ii) hypoxia. We found that ischemia, but not SD alone, caused a delayed depolarization resulting from the replacement of intracellular K^+^ with Na^+^, and also produced secondary changes to Ca^2+^ and Cl^−^. We also found spatial effects, with increased vulnerability to ischemia-induced excitotoxicity (from Ca^2+^) for basal dendrites, contrasting with greater susceptibility to dendritic beading (from Cl^−^) for apical dendrites. These distinctive patterns of flux are likely not only to lead to different levels of risk of cell-death but also to different types of cell damage, which will have different consequences for cell and network dysfunction. Different therapeutic strategies targeting these distinct mechanisms may thus be optimized to mitigate the kinds of damage that predominate under the conditions of hypoxia versus normoxic SD.

## 2 Methods and Materials

Our model is based on previous work utilizing a data-driven simulation workflow to create a hippocampal CA1 pyramidal neuron model with detailed morphology and appropriate mechanisms and parameters to provide a good fit to experimental recordings (ModelDB:244688) (Migliore et al., 2018). We chose CA1 as it has been extensively studied in ischemia and SD(Silver and Erecińska, 1990; Pérez-Pinzón et al., 1995; Larsen et al., 2005; Bennett et al., 2023), the large pyramidal neurons are particularly vulnerable to metabolic challenges(Bartsch et al., 2015) The model includes two intracellular shells with radial diffusion between the two, one (between 0.85 of the radii and the plasma membrane) representing surface concentrations and one (from the center to 0.85 of the radii), which in this model is further subdivided into the cytosol (83% of the volume) and the endoplasmic reticulum (ER) (17% of the volume). The concentrations in the surface and cytosol are closely coupled by diffusion. Subscripts are used to indicate concentrations in the different regions: *i* for the surface regions, *cyt* for cytosolic, and *o* for extracellular.

The model includes both intracellular and extracellular reaction-diffusion, using the NEURON simulator (Hines et al., 2019; Carnevale and Hines, 2006) including its reaction-diffusion module, which supports coarse-grained volume-averaged models of the extracellular space (McDougal et al., 2013; Newton et al., 2018).

### 2.1 Energy and metabolism

We explicitly modeled oxygen and ATP concentrations: Oxygen is consumed to produce ATP and restored at a rate proportional to how depleted it is relative to a baseline concentration for normoxic or ischemic/hypoxic tissue Wei et al. (2014).

We used a simplified version of the ATP model Le Masson et al. (2014), where ATP returns to a set steady-state (ATP_*ss*_) of 2.59 mM (Veech et al., 1979) with time constant 3.9 ms. This process depends on the amount of O_2_ available, with 6 O_2_ molecules capable of producing 30 molecules of ATP (Rich, 2003). The model includes rapid intracellular diffusion of ATP, and both intracellular and extracellular diffusion of O_2_ (Hubley et al., 1996; Gerkau et al., 2019). For simplicity, rates of the energy-dependent Na^+^-K^+^ pump (Noske et al., 2010), SERCA (Mahmmoud, 2008), and the plasma membrane calcium pump (PMCA) were modified to include Michaelis-Menten kinetics for their dependence on ATP, while keeping the same maximum rate.

### 2.2 Glutamate and synapse models

Our homeostatic glutamate model was based on Hübel et al. (2017), where each segment is equipped with an “average-synaptic cleft” and glutamate release is based on a threshold of the membrane potential of the soma and the available stores. We model the neuron by dividing the morphology into 507 compartments called segments in NEURON (by limiting the maximum length of a compartment 40 μm). We equip each of these compartments with a single “average-synaptic cleft” model. This simplification allows us to model excess glutamatergic drive that occurs during spreading depolarization without modeling each of the ∼ 30, 000 synapses (Megías et al., 2001). Glutamate is released into synapse for the *i*^th^ segment at a rate *J*_*i*_ when the membrane potential passes a given threshold *v*_*crit*_.

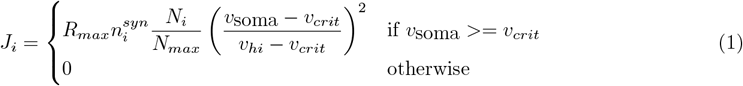

Here *N*_*max*_ is the maximum amount (fmol) of stored glutamate and *N*_*i*_ is the current store for the *i*th section. *R*_*max*_ is the maximum release rate, *v*_*crit*_ and *v*_*hi*_ are the range of membrane potentials where glutamate is released, and 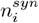 is the number of synapses on the segment. Release of glutamate is also simplified, it only depends on the somatic membrane potential.

Uptake occurs from both the cleft and the extracellular space (ECS) and follows Michaelis-Menten kinetics. Glutamate can diffuse between the ECS and the synapses. Glutamate is recycled, replenishing the stored glutamate and (*N*_*i*_). N-methyl-D-aspartate receptors (NMDAR) and α-amino-3-hydroxy-5-methyl-4-isoxazolepropionic acid receptors (AMPAR) were also adapted from Hübel et al. (2017), with NMDAR Ca^2+^ currents based on Somjen et al. (2008).

### 2.3 Extracellular concentrations and clearance

We simulated diffusion of molecules (K^+^, Na^+^, Cl^−^, Ca^2+^, O_2_ and glutamate) in the extracellular space bound by the neuron’s convex hull, as approximated with Delaunay tessellation (Virtanen et al., 2020). The hull is required to avoid modeling an unrealistically large empty volume of ECS. The effective free volume was reduced by a factor of 5, since multiple neurons would pass through this region, but we were only modeling a single neuron within this space. Our simulation of the extracellular space utilizes Neumann (zero flux) boundary conditions, (so K^+^ does not diffuse out of the model). This boundary models the situation where the neuron and glia sit within a larger volume of tissue experiencing identical pathological conditions.

An astrocyte model based on Kager et al. (2002); Somjen et al. (2008) with a simplified morphology is included. The same homeostatic mechanisms that feature in the neuron model are included in the astrocyte.

### 2.4 Calcium dynamics

The original model (Migliore et al., 2018) included simple calcium decay in the two radial shells. It used different parameters for the soma, axon, apical dendrites, and basal dendrites, with the *I*_h_ and K_*A*_ conductances and the passive leak potential dependent on their distance from the soma. Here, the passive leak was replaced with ion-specific leaks independent of distance from the soma.

We expanded this model to include mechanisms supporting calcium-induced-calcium release from (Neymotin et al., 2016) (ModelDB:185858): a calcium buffer, IP_3_ receptors, Ryanodine receptors, and a simplified pathway that increases IP_3_ concentrations in response to metabotropic glutamate receptor (mGluR) activation. We simplified the mGluR mechanisms; rather than modeling the signaling pathway, we only included the essential feature that IP_3_ increases in response to glutamate binding to mGluR.

The Na^+^-Ca^2+^ exchange is an important mechanism in maintaining homeostasis and is implicated in excitotoxicity. Here, we use a ping-pong bi-bi cyclic model and parameters from the mouse cardiac myocyte (Weber et al., 2001) as the NCX1 from of the Na^+^-Ca^2+^ exchange is expressed both in the heart and brain. The model has a 3:1 stoichiometry and reversal potential of −53 mV (with the initial ion concentrations), closer to the resting membrane potential (RMP) of the neuron −69.5 mV, than the alternatives. The Na^+^-Ca^2+^ exchange model has parameters for saturation with each intracellular and extracellular ion, making this scheme more appropriate for simulating pathology. The density of the Na^+^-Ca^2+^ exchange or its maximum velocity was determined by parameter optimization.

### 2.5 Distance dependent mechanisms

A-type channel (KA) and the HCN *I*_h_channel conductances are known to increase with distance from the soma in pyramidal cellsMigliore and Shepherd (2002). We have incorporated this phenomenon into the model by having *I*_h_ channel conductance increase linearly with slope 0.03 and KA conductance governed by a sigmoid with length scale 50 μm based on the path distance to the center of the soma. While cAMP is a potent agonist of the HCN channel and the Na^+^-K^+^ pump, it is likely to decrease during ischemia as the majority of ATP will be utilized by ATPases, rather than catalyzed to cAMP. The ATPases modeled are the; Na^+^-K^+^ pump, plasma membrane Ca^2+^ ATPase (PMCA) and sarcoendoplasmic reticulum Ca^2+^ transport ATPase (SERCA).

The *I*_h_ is permeable to sodium and potassium with a relative permeability of 0.36 Magee (1998); Aponte et al. (2006). We used this and the Goldman–Hodgkin–Katz (GHK) equation to divide the current between the two cations while keeping the same gating dynamics as the original model Migliore et al. (2018).

### 2.6 Parameter fitting

Following the protocols in Migliore et al. (2018) the model parameters were optimized to reproduce 12 electrophysiological features and ensure ionic homeostasis in the soma, an apical, and a basal section in response to two stimiuli cases. To keep fits physiologically plausible, any tests with negative conductance values were assigned the worst possible fitness score; no simulations were run in this case. To speed up parameter optimization, the three mechanisms that alter the intracellular chloride concentration (NKCC1, KCC2, and the ion-specific leak) were combined to ensure a non-negative leak conductance for each parameter set.

The optimizations were performed using BluePyOpt (Van Geit et al., 2016) with over 600 generations, with a population of 256 parameter sets per generation over 150, 000 simulations in total, each taking ∼ 23 minutes for each second of simulated time on a single core of an Intel Xeon E5.

In (Migliore et al., 2018), parameters for the apical dendrites, basal dendrites, and the soma were allowed to vary independently. In this work, to understand the role of morphology and reduce the search space for plausible models, most ion channel conductances were the same between the apical and the basal dendrites. The exceptions were *I*_h_ and k_*A*_, which depended on the distance from the soma (Magee, 1998), the glutamate mechanism, NMDAR, and AMPAR, which were more densely distributed in the apical dendrites (Megías et al., 2001).

The full model code is freely available on ModelDB (McDougal et al., 2017); https://modeldb.science/2017004 (access code: review)

### 2.7 Human tissue neuropathologic evaluation

Deidentified archival tissue blocks of adult human brain from the Yale Department of Pathology were reviewed by a neuropathologist and sections containing subacute infarct were selected for further workup. Sections were stained using immunohistochemistry with a primary antibody against neurofilament (Agilent Clone 2F11, Catalog #: GA60761-2) and hematoxylin counterstain. Slides were imaged on a Motic EasyScan Infinity 60 imaging system (Motic Inc., Kowloon, Hong Kong) at 40X magnification, at 0.26 μm/pixel resolution. The resulting images were reviewed in QuPath (Bankhead et al., 2017).

## 3 Results

Neuropathologic review of post-mortem human brain tissue from three subjects of ages 62-75 years with subacute infarcts of cerebral cortex was performed using immunohistochemistry for neurofilament (abundant heteropolymer neural proteins that contribute to maintenance of neuronal structure). Beading, characterized by periodic varicosities (i.e. swollen foci containing neurofilament) along the length of neurites was observed in both the penumbra and within the ischemic core of the infarcted tissue, but not in adjacent uninvolved cortex (Figure 1).

**Figure 1.**
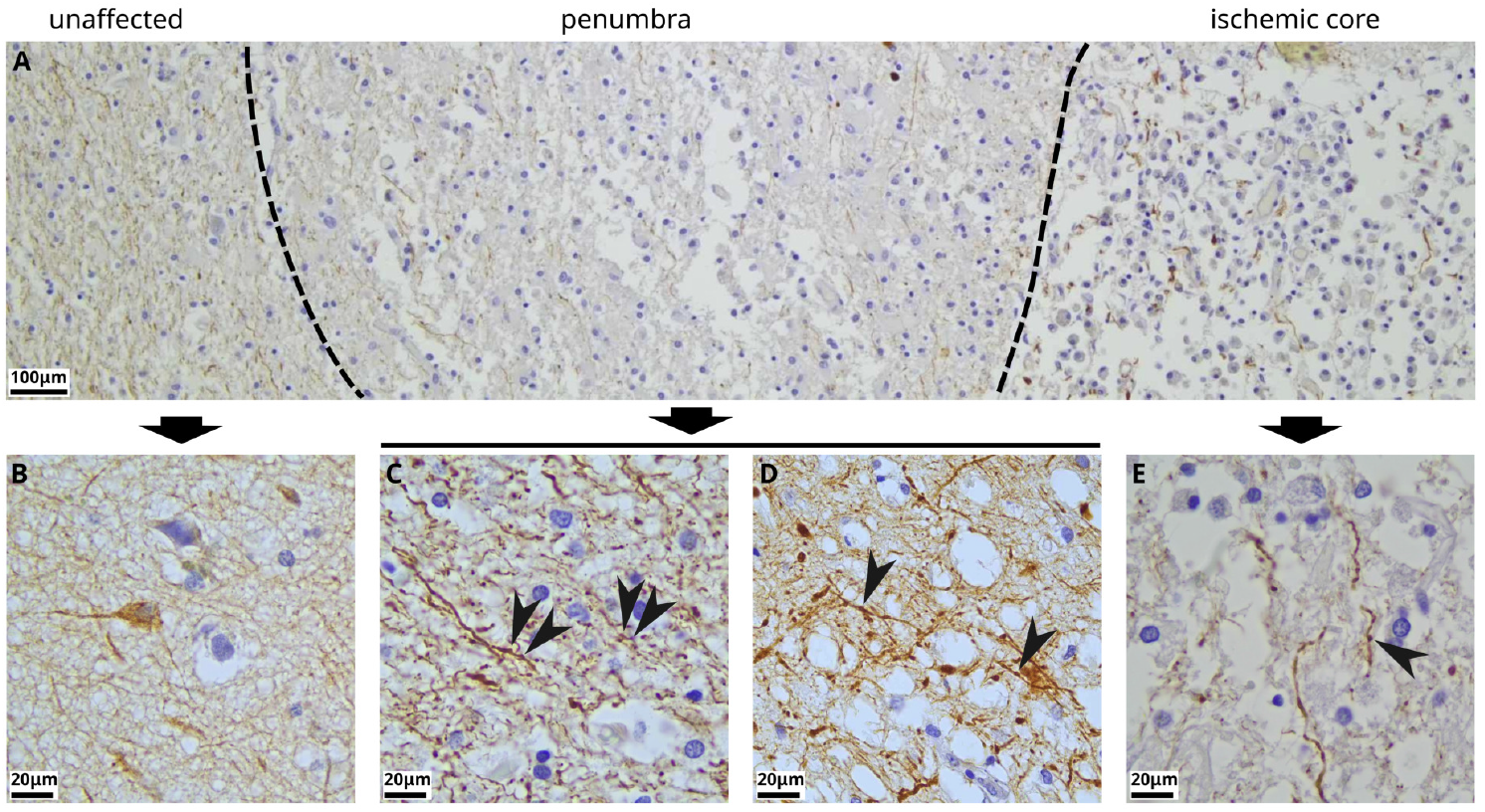
An example of neurite varicosities (“beading”) in the penumbra and the core of human cortex infarct, as shown with immunohistochemistry for neurofilament. **(A)** Low-power view demonstrating the zones of brain tissue affected by subacute infarct. At left, in a region of the gray-white matter junction (with occasional layer VI neurons), that is relatively spared of ischemic injury, there is more organized neuropil occupied by neurons, oligodendroglia, and reactive astrocytes. In the penumbra of the infarct, there is vacuolization of the neuropil and infiltration by macrophages, with some loss of tissue integrity. In the ischemic core, there is severe loss of brain tissue, with cavitation and near-replacement of neuropil by macrophages, and only disorganized and dissociated neurites surviving, with prominent swelling. **(B)** Enlarged view of unaffected tissue with no visible neurite varicosities. **(C**,**D)** Enlarged view of penumbra with arrowheads indicating examples of neurite beading. **(E)** Enlarged view of the ischemic core with arrowheads indicating examples of neurite beading.

Because histologic sections are thin (5*μ*m) slices and thus only show small parts of the dendritic arbor, and we sought a more comprehensive understanding of 1) the mechanisms of SD and hypoxic SD (‘HSD’) in humans, 2) the extent to which they drive ischemic damage, and 3) how neurite morphology contributes to vulnerability, we developed multiscale 3-dimensional models of CA1 pyramidal neurons with simulated SD and HSD. We hypothesized that the high density of dendrites near the soma would give rise to different vulnerabilities in different parts of the dendritic arbor under these conditions. In particular, we hypothesized that the densely packed basal dendrites would be more vulnerable due to positive feedback loops from other dendrites where increased extracellular K^+^ or glutamate would cause neighboring dendrites to depolarize, releasing more K^+^, while also increasing the demand on the Na^+^-K^+^ pump consuming more ATP reducing the ability for the dendrite to respond to further stress.

### 3.1 Membrane potential changes under hypoxia were slower but larger than those due to elevated extracellular potassium

Under physiological conditions, neurons respond to moderate depolarizing stressors with a return to baseline resting membrane potential (RMP; at the end of the burstFigure 2). Following a common experimental protocol (Hamill et al., 1981; Perkins, 2006), we depolarized the neuron with a 500 ms current injection at 0.5 nA. This current produced a large burst of APs resulting in large ionic fluxes, with the influx of Na^+^ and efflux of K^+^. Homeostasis was quickly restored in all compartments, as pumps quickly restored ion concentrations and recovered Nernst potentials. By their nature and by the distribution of ions, even when the cell is at rest, sodium channels admit a small amount of sodium into the cell and release a small of potassium to the extracellular space. Under normoxic conditions, pumps compensate for these ion fluxes, and the concentrations and corresponding Nernst potentials are held approximately constant (dashed lines in Figure 2a). Following current injection, there was little change in V_m_ throughout the neuron for the rest of the simulation (Figure 2b).

**Figure 2.**
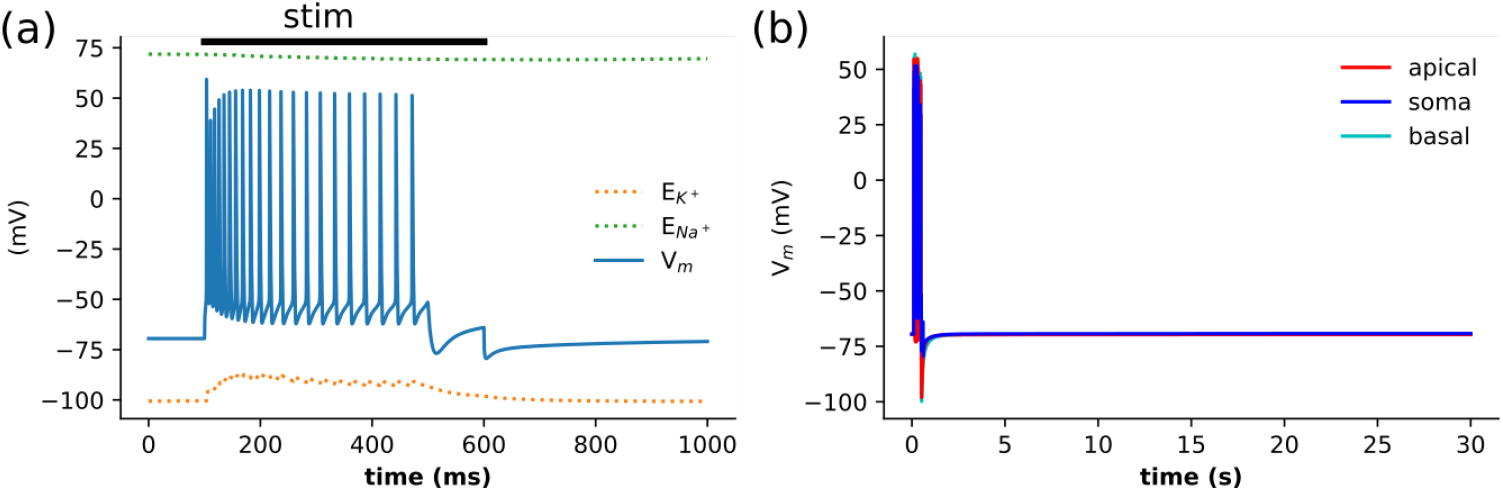
Control: membrane potential returns to baseline following 500 ms somatic current stimulation. **(a)** The first 0.5 s of somatic membrane potentials shows a burst of APs followed by depolar-ization block and quick return to RMP at the end of the stimulus. Nernst potentials remain relatively static, with a small rise in E_K_ from [K^+^]_o_ changes due to APs. **(b)** Following the burst of APs that depolarized all neuronal compartments, membrane homeostasis was rapidly restored and maintained indefinitely – 1 min of simulation shown.

Elevated extracellular K^+^ rapidly produced a sustained depolarization with variation on V_m_ across the dendritic tree, since depolarization occurred throughout the dendritic tree, eliminating the dendrites as current sinks (Figure 3a-c). (1) We ramped up extracellular K^+^ to 10 mM in the first 1 ms; (Figure 3b) (2) E_K_ followed K^+^ passing V_m_, (3) and thereby reducing and even briefly reversing *I*_K_ to an inward current (4). This depolarized the cell towards the spike threshold (5) producing the sudden downward (inward) *I*_Na_ deflection (where Na^+^ *m*_∞_ hits inflection) (6) triggering a single AP. (7) Incomplete repolarization to a sustained depolarization was consistent with SD (Figure 3c). (8) During the sustained depolarization, there were relatively small changes in V_m_, which were different for different parts of the neuron. Notably, a decline in apical V_m_, as there was less competition for available O_2_ in the apical than in the dense dendritic arbor near the soma.

**Figure 3.**
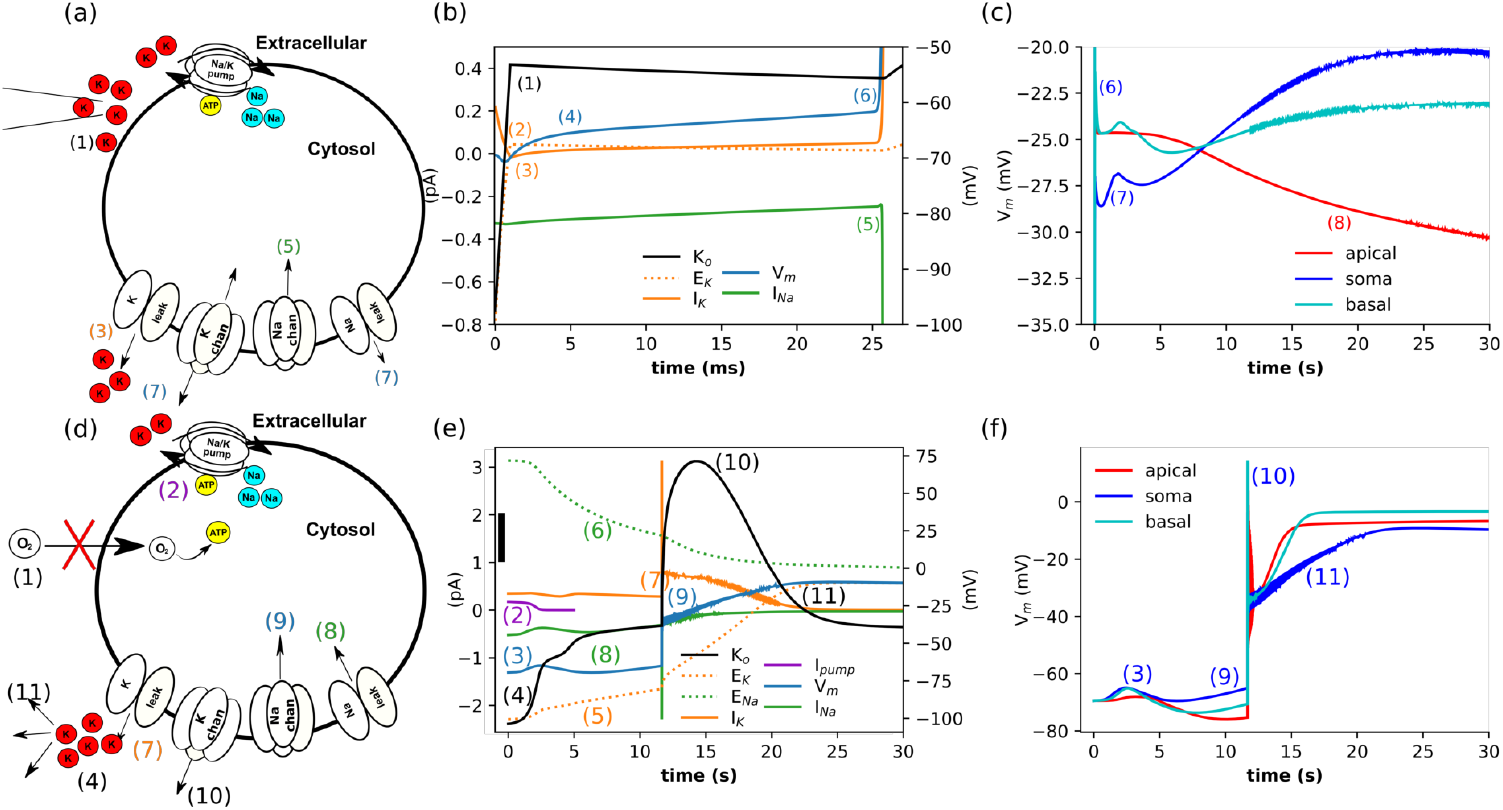
Either elevated extracellular K^+^ or hypoxia gave rise to sustained depolarizations, ↑[K^+^]_o_ produced rapid depolarizations, whereas hypoxia led to slower but larger depolarizations. **(a-c)** Elevated extracellular K^+^ causes a spike followed by sustained depolarization. Sequence of events (numbers on panels): (1) elevation of extracellular K^+^ from 2.9 mM to 10 mM; (2) E_K_ increase above the V_m_; (3) *I*_K_ leak reverses; (4) neuron depolarizes; (5) V_m_ approaches spike threshold (∼ −60 mV); (6) producing the upswing of AP; (7) at the end of the spike, continued Na^+^ currents create a new equilibrium. (8) apical V_m_ falls while somatic and basal rise, with small high frequency oscillation. **(a)** Schematic. **(b)** Currents *I*_K_ and *I*_Na_ are shown the left axis, potentials E_K_ and V_m_ on the right axis. Changes occurred rapidly, on a millisecond timescale. **(c)** A single AP (truncated y-axis) is followed by incomplete repolarization and sustained depolarization consistent with SD. Regional variations in V_m_ result from extracellular differences in O_2_ and K^+^ arising from the amplifying effects of interacting dendrites near the soma. Concentrations in a compartment of the basal dendrites 201 μm, and apical dendrites 611 μm from the soma are shown for comparison. **(d-f)** Another route to SD: hypoxia reduces pump activity and ion homeostasis fails. Sequence of events: (1) ↓O_2_ →↓ATP; (2) ↓Na^+^-K^+^ pump; (3) ↑V_m_; (4) ↑[K^+^]_o_; (5) ↑E_K_; (6) ↓E_Na_; (7) ↓*I*_K_; (8) ↓*I*_Na_; (9) V_m_ approaches spike threshold producing a partial AP; (10) large *I*_K_ led to greater accumulation extracellular K^+^ and further sustained depolarization with small high-frequency oscillations; (11) excess extracellular K^+^ diffuses away as the neuron and V_m_ remains depolarized as K^+^ is replaced with Na^+^. **(d)** Schematic. **(e)** Currents *I*_K_ and *I*_Na_ are shown on the left axis, excluding the contribution of Na^+^-K^+^ pump; potentials E_K_, E_Na_, and V_m_ on the right axis; [K^+^]_o_ 2.9 to 5.9; scale bar 0.5 mM. **(f)** The variation in V_m_ between the soma and the dendrites. A compartment of the basal dendrites 201 μm, and apical dendrites 611 μm from the soma, are shown for comparison.

By contrast, sustained hypoxia resulted in delayed depolarization (Figure 3e-f). A relatively small initial change in V_m_ (3) masked substantial shifts in Nernst potentials: E_Na_ (6) dropped, while E_K_ rise (5). The hypoxic-induced delayed depolarizations can be viewed as a sequence of events: (1) rapid reduction of O_2_ supply (hypoxia). (2) Na^+^-K^+^ pump current fell to 1% of its initial magnitude over 2.95 s as ATP consumed (Figure 3e); (3) ↑V_m_ rose with loss of outward Na^+^-K^+^ pump current; then fell due to K^+^ channel responses to V_m_. (4) ↑[K^+^]_o_ due to Na^+^-K^+^ pump failure, (5) ↑E_K_, (6) ↓E_Na_, (7) reduced outward *I*_K_ was less than (8) reduction in inward *I*_Na_; (9) elevated V_m_ to spike threshold, which produced only a partial AP due to the fall in E_Na_. Despite inactivation of the Na^+^ channel shortly after opening (falling below 1% of peak conductance by 15 s), the neuron cannot repolarize due to the elevated E_K_. (10) ↑ [K^+^]_o_ and further depolarization, produced by large and sustained *I*_K_. (11) ↓[K^+^]_o_ due to extracellular diffusion, while V_m_ remained elevated as intracellular K^+^ continued to be replaced with Na^+^. Small high-frequency oscillations in the V_m_ at the soma were produced by Na^+^ channel activity. Changes in extracellular concentration were smaller than intracellular changes as the size of the extracellular space exceeds that of the neuron, even with the currents scaled 5-fold (see methods).

During sustained hypoxia, synaptic mechanisms produced distinct somatic and dendritic responses (Figure 3f). The sequence of events that affected the somatic membrane potential produced similar disruptions in the dendrites: (3) ↑V_m_ was similar at the soma and basal dendrites, as Na^+^-K^+^ pump fell rapidly (to less than 1% at the soma and ∼ 6% at the basal dendrites within 3 s) with many neighboring dendrites competing for limited O_2_. In contrast, the sparser apical dendrites were relatively spared, with little change in the Na^+^-K^+^ pump within 3 s, eventually reaching a plateau of 62%. (9) The range of membrane potentials also results from the residual Na^+^-K^+^ pump current, which was largest in the apical dendrites. (10) The partial AP triggered glutamate release, which activated synaptic mechanisms in the dendrites that were absent in the soma (particularly AMPAR). These synaptic mechanisms produced a larger and more rapid rise in dendritic V_m_ than somatic V_m_, despite similar extracellular environments for the soma and basal dendrites. (11) ↓[K^+^]_o_ due to extracellular diffusion, and was far higher around the soma due to efflux from the soma and multiple neighboring dendrites. However, V_m_ remains elevated due to the almost complete loss of [K^+^]_i_, with E_K_ ∼−5 mV. Compared to elevated extracellular K^+^ (Figure 3c) where glutamate was also released by the initial AP. However, the synaptic currents that led to higher dendritic than somatic V_m_ in Figure 3f, were dwarfed by greater changes caused by the large and rapid rise of E_K_.

Sustained hypoxic effects were delayed by seconds rather than the millisecond shifts seen with high K^+^, reflecting the temporal buffering provided by the time required for ATP consumption and pump failure (compare Figure 3d-f with Figure 3a-c). However, depolarization, once it occurred, was considerably more pronounced under hypoxia as without ATP the pumps were unable to compensate. High extracellular K^+^ and hypoxia both led to the breakdown of ion homeostasis, but over different timescales and via distinct mechanisms. The rapid initial increase in extracellular K^+^ can be viewed as the loss of charge from one of the neuron’s batteries (E_K_), which led to a rapid initial disruption of ion homeostasis (initial 25 ms shown in Figure 3b). The loss is followed by a slower secondary rise in E_K_ and a fall in E_Na_ further reducing the neuron’s batteries as V_m_ rises. In contrast, the rapid reduction in O_2_ produced slower initial changes in both Nernst potentials and V_m_ (Figure 3e). After ∼ 12 s, the partial AP led to the almost complete loss of E_K_ and E_Na_, producing much larger shifts in V_m_. These differences between hypoxic and elevated [K^+^]_o_ conditions are partly due to the Na^+^-K^+^ pump, a current source that charges the neuron’s batteries. Hypoxia limited the capacity of the Na^+^-K^+^ pump and elevated K^+^ increased the drive; in both cases, the drive exceeded capacity. However, the residual Na^+^-K^+^ pump current at the soma was 9.8 fold greater under elevated K^+^ than when O_2_ was reduced.

Hypoxia and elevated K^+^ occur in hypoxic spreading depolarization (HSD) in the penumbra of ischemic stroke. Simulation of both insults together showed the initial dynamics are dominated by the effects of elevated K^+^, similar to Figure 3b. Over time, the membrane potentials were similar to those for hypoxic conditions (Figure 3f), resulting from the breakdown of ion homeostasis.

### 3.2 Ischemia and SD produced different distributions of ion concentration in the neuron

Disruption of ion homeostasis caused highly variable Ca^2+^ accumulation across neuronal compartments, revealing compartment-specific vulnerabilities driven by local energy supply, transporter activity, and morphology. (Figure 4). Elevated extracellular K^+^ (a model for SD) led to orders-of-magnitude increase in Ca^2+^ concentrations throughout, from 60 nM into the mM range (Figure 4a,c). The Ca^2+^ increase was greatest in the basal dendrites, with comparable values in some proximal apical dendrites, while distal apical dendrites showed less accumulation. Some dendritic compartments were spared significant rises in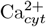. The large range in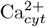 concentrations reflects a pattern of vulnerabilities in the neuron. Different patterns of NMDAR and Na^+^-Ca^2+^ exchange currents determined the differences in Ca^2+^ in basal vs. apical dendrites. Tightly packed basal dendrites competed for limited O_2_ and suffered greater reduction in ATP than the distal apical dendrites. Residual Na^+^-K^+^ pump activity was lower in the basal dendrites, which led to higher [Na^+^]_i_, and thus limited Ca^2+^ clearance via the Na^+^-Ca^2+^ exchange (Figure 4(e),(g)) Further reduction in Ca^2+^ elimination due to decreased ATP-limited Ca^2+^ pump and SERCA activity. NMDAR activity depended on accumulation of glutamate, while Na^+^-Ca^2+^ exchange increased with elevated [Na^+^]_i_ and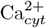. The rise in [Na^+^]_i_ was most pronounced in compartments where the Na^+^-K^+^ pump was overwhelmed and ATP declined. The role of both ATP and glutamate tied these changes to the morphology.

**Figure 4.**
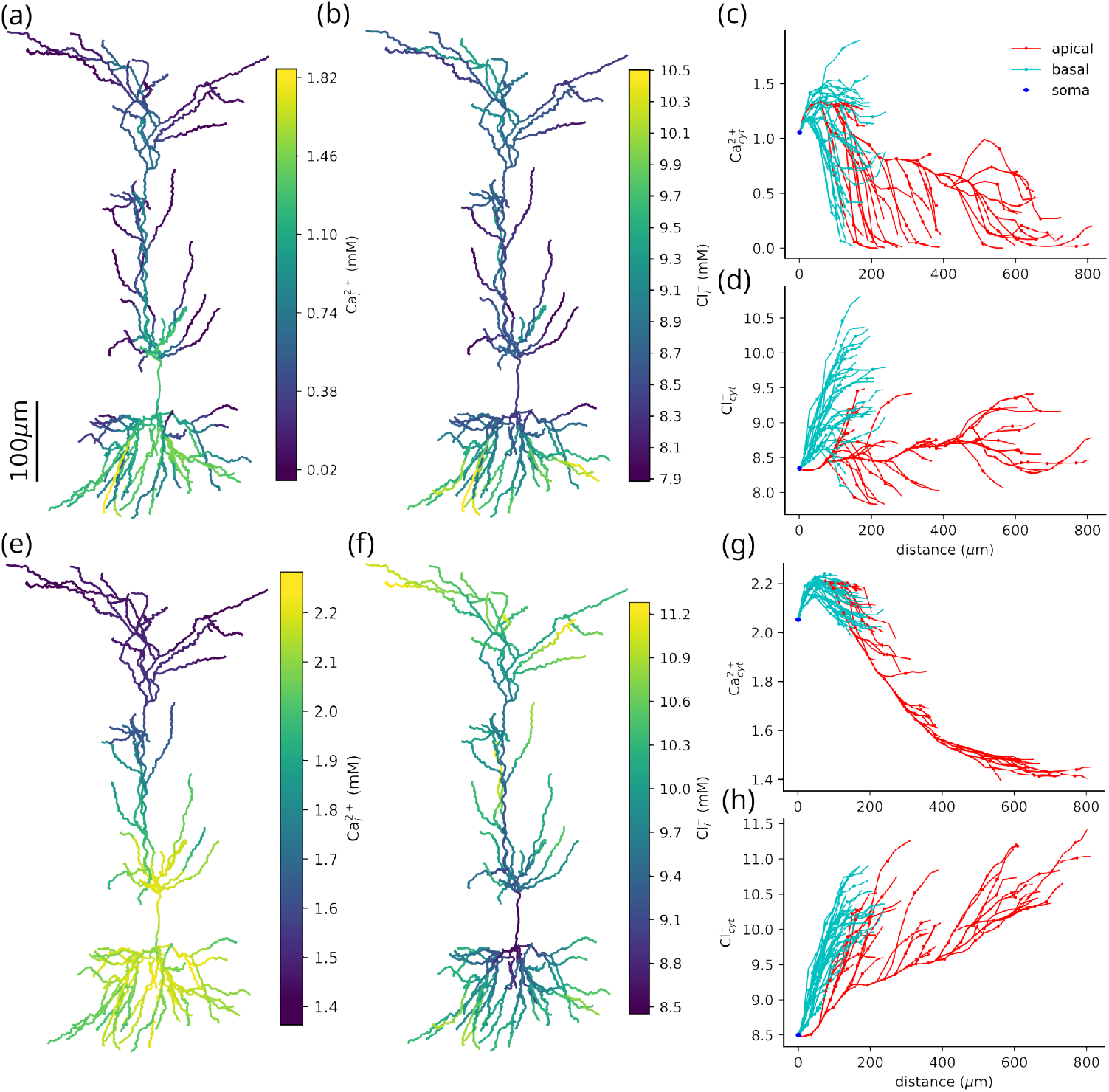
Elevated K^+^ and hypoxia produce different distributions of Ca^2+^ and Cl^−^. Both Ca^2+^ and Cl^−^ are most elevated in the basal dendrites under normoxic conditions with elevated K^+^(a-d). Hypoxia induced elevated Ca^2+^ in the basal dendrites, with greater accumulation of Cl^−^ in the distal apical dendrites (e-h). Under normoxic conditions with elevated K^+^: **(a)**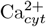 differed in soma (1.06 mM), basal dendrites (1.20 mM to 1.10 mM), apical dendrites (1.31 mM to 0.11 mM). **(b)**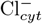 increased with distance from the soma (8.35 mM) in the basal dendrites (8.38 mM to 9.77 mM), whereas apical dendrites have a lower concentration that does not depend on distance from the soma (8.32 mM to 8.64 mM). **(c)**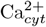 peaked in the basal dendrites (at 1.90 mM 176 μm from the soma) **(d)**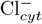 also peaked with Ca in the basal dendrites (at 9.49 mM 176 μm from the soma). Under hypoxic conditions: **(e)**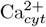 was most elevated at the soma (2.05 mM) and basal dendrites (2.07 mM to 2.01 mM), with similar concentrations in the proximal apical dendrites and lower concentrations in the distal apical dendrites (2.18 mM to 1.44 mM). **(f)**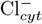 increased with distance from the soma (8.50 mM in both basal (8.53 mM to 10.53 mM) and apical dendrites (8.51 mM to 11.02 mM), **(g)**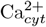 peaked in the basal dendrites (2.24 mM at 82 μm from the soma) and declined in the apical dendrites with distance from the soma. **(h)**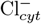 accumulation increased linearly with distance from the soma 1.60mM/mm (R^2^ = 0.368), with greater rises in the apical 2.19mM/mm (R^2^=0.75) than basal 1.03mM/mm (R^2^=0.57) dendrites. Concentrations ranges in the dendrites were taken at the closest and furthest parts from the soma; apical dendrites (154 μm and 773 μm) and basal dendrites (35 μm and 203 μm).

The rise of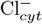 was less marked, increasing up to 48% from a baseline of 6.6 mM (Figure 4(b)&(d)). The pattern of Cl^−^ accumulation under elevated K^+^, was similar to that of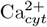. Concentrations of Cl^−^ were higher in the basal than the apical dendrites. The reversal of KCC2 drove the Cl^−^ increase as the dominant Cl^−^ current. KCC2 normally utilizes the K^+^ gradient to remove Cl^−^ from the neuron, but may reverse direction under pathological conditions. KCC2 reversal occurred almost immediately, due to the shift in E_K_ that resulting from elevated extracellular K^+^ (Figure 3b). In parts of dendrites KCC2 recovered, as there was sufficient ATP to maintain some Na^+^-K^+^ pump activity, increasing [K^+^]_i_ and restoring E_K_ above E_Cl_.

Under hypoxic conditions, the rise in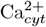 was more pronounced with greater differences between proximal and distal dendrites (Figure 4(e)&(g)). As with the normoxic case, the rise in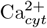 depended on the balance between NMDAR and Na^+^-Ca^2+^ exchange, which tie Ca^2+^ concentrations to glutamate and O_2_, which are affected by the dense dendritic arbor near the soma.

The distribution of Cl^−^ was well-described by path distance from the soma, with the increase more pronounced in apical dendrites. Concentrations of Cl^−^ rose with distance from the soma at 1.60 mM/mm in our simulation, with slightly greater increases with distance in the basal than in the apical dendrites. Unlike under elevated K^+^, under hypoxic conditions, KCC2 reversed throughout the neuron, with differences in Cl^−^ caused by the timing and extent of the reversal. KCC2 reversal was delayed as the changes in E_K_ were slower under hypoxic conditions (Figure 3e). KCC2 reversal occurred first in a basal dendrite before the apical dendrites. The greater increase in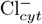 in the apical dendrites resulted from the breakdown of the E_K_ and E_Na_, which were both lower on average in the apical dendrites. KCC2 is driven by the difference between E_K_ and E_Cl_, and NKCC1 (a sodium-potassium-chloride transporter) by the difference between both E_K_ and E_Na_ with E_Cl_. Changes in Cl^−^concentration distribution were driven primarily by the non-uniform disruption to K^+^ and Na^+^. Basal dendrites near the soma experience higher [K^+^]_o_ than apical dendrites (Figure 6c). As a result, E_K_ was higher near the soma, so there was greater KCC2 activity, consequently less elevated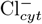.

In both cases, energy-dependent clearance mechanisms could better respond to demand in apical compared to basal dendrites. Glutamate-dependent NMDAR activation significantly increased Ca^2+^ accumulation and altered depolarization dynamics. We identified the contribution of glutamate-NMDAR influx by assessing a model without NMDAR. Under hypoxic conditions, this model produced a similar distribution of Ca^2+^ and Cl^−^ with lower overall intracellular concentrations. In the absence of glutamate elevated K^+^ led to large high frequency oscillations rather than sustained depolarizations and resulted in a far smaller rise in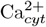.

### 3.3 Hypoxia contributes to pathological changes of calcium and chloride throughout the cell

Elevated extracellular K^+^ led to immediate rises in Ca^2+^ and Cl^−^ (Figure 5a-c), that varied with location and available O_2_. Under hypoxic conditions,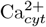 rose most rapidly in the apical dendrites before it plateaued and was surpassed by the basal and somatic concentration Figure 5a solid lines. Under normoxic conditions, elevated [K^+^]_o_ produced substantial rises in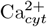 at the basal dendrites and soma. In contrast, apical dendrites were able to maintain lower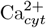 due to less competition for O_2_ and ATP Figure 5a dashed lines. The ER Ca^2+^ initially declined, but then reversed and recovered as it reached equilibrium with the elevated cytosolic concentration Figure 5b. Again, apical dendrites were spared in normoxic conditions. The rise in Cl^−^ was most pronounced in the apical dendrites, for both normoxic and hypoxic conditions Figure 5c. There was less competition for available O_2_ in apical dendrites compared with the densely packed basal dendrites, which led to higher [K^+^]_i_ and lower E_K_, leading to lower KCC2 Cl^−^influx or recover KCC2 Cl^−^efflux in some dendrites under normoxic conditions.

**Figure 5.**
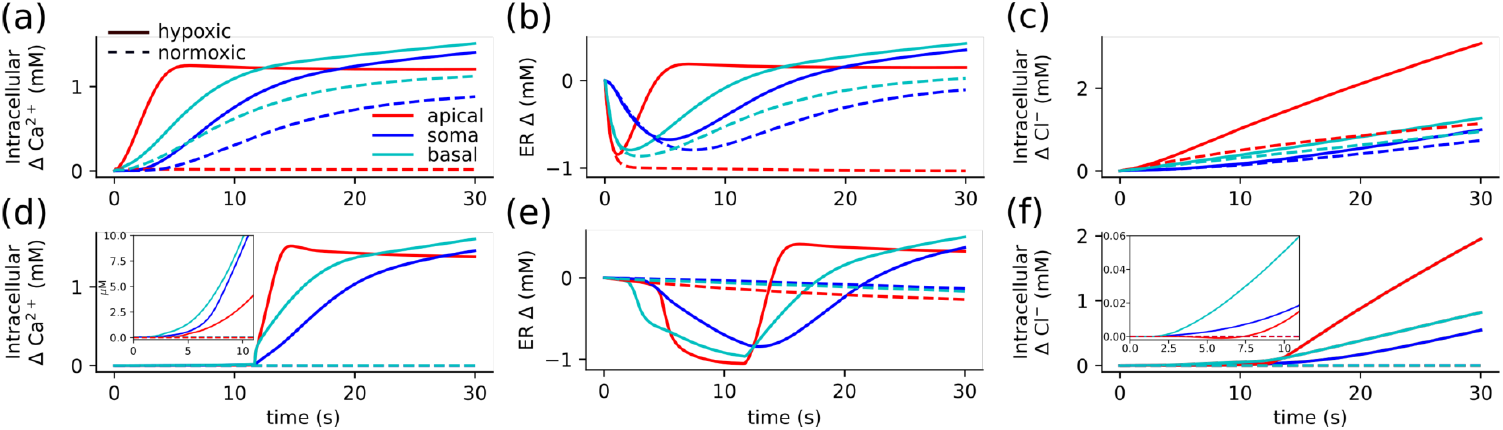
Elevated K^+^ led to a rapid rise in both the Ca^2+^ and Cl^−^ concentrations, while hypoxic changes are delayed. Rapid concentration changes followed neuronal depolarization that occurred almost immediately (∼ 25 ms) with elevated K^+^ (a-c), or (∼ 12 s) with baseline K^+^ under hypoxic conditions (d-f). **(a)** Under hypoxic conditions (solid lines), the rise in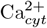 was most pronounced in the apical dendrites, whereas under normoxic conditions, the apical dendrites were spared. **(b)** Ca^2+^ in the ER showed a decline followed by recovery as it equilibrates to the elevated intracellular Ca^2+^. **(c)** Cl^−^ concentration rose gradually in the soma and basal dendrites and had the greatest increase in the apical dendrites under both hypoxic and normoxic conditions. In the absence of applied K^+^ (d-f) there was little change in concentrations under normoxic conditions (dashed-line), with hypoxia leading to substantial changes following depolarization at ∼ 12 s. **(d)** The smaller initial increase in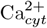 were due to influx from the ER and brief reversal of Na^+^-Ca^2+^ exchange (between 3.08 s and 4.16 s at the soma). The rise in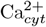 following the AP was initially greatest in the apical dendrites, which then plateaued. Whole basal and somatic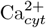 rose more slowly; they ultimately exceeded apical concentrations. **(e)** The decline in ER Ca^2+^ preceded the depolarization, which reversed following depolarization as ER Ca^2+^ rose to equilibrate with cytosolic concentrations. **(f)** After an initially rapid rise in the basal dendrites (inset), the long-term rise in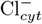 was greatest in the apical dendrites.

With baseline K^+^ under hypoxic conditions, disruption of the balance between K^+^ and Na^+^ produced pathological changes in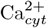 and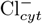 Figure 5d-f. The majority of the pathological accumulation of Ca^2+^ and Cl^−^ occurs with depolarization at ∼ 11 s. Prior to depolarization, cytosolic Ca^2+^ increased gradually throughout the neuron by about two orders of magnitude, from 60 nM to μM concentrations. These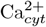 dynamics were driven by the ER store, Na^+^-Ca^2+^ exchange, and buffering. Somatic Ca^2+^ rose substantially pre-depolarization, (Figure 5d inset) aided in part by a brief reversal of the Na^+^-Ca^2+^ exchange. Basal dendrites showed a comparable rise, while the apical concentration rose about a third as much. The Na^+^-Ca^2+^ exchange clearance was reduced and only briefly reversed in parts of the dendrite and at the soma. In all regions, ER Ca^2+^ was largely depleted during the pre-depolarization phase (Figure 5e solid line). This decrease exceeds the increase in cytosolic Ca^2+^ even after adjusting for differences in volumes, likely due to cytosolic Ca^2+^ buffering and residual clearance via Na^+^-Ca^2+^ exchange and Ca^2+^ pump.

Depolarization at ∼ 11 s triggered a rapid rise in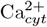 driven by NMDAR and modulated by Na^+^-Ca^2+^ exchange, leading to compartment specific dynamics Ca^2+^ changes (Figure 5d solid line). Dendritic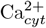 increased by almost two orders of magnitude in less than two seconds. Apical Ca^2+^ rose most rapidly, followed by basal Ca^2+^, however, apical concentrations plateaued. Dendritic concentrations were driven by the balance between NMDAR Ca^2+^ influx and Na^+^-Ca^2+^ exchange efflux. After the rapid rise, the Na^+^-Ca^2+^ exchange became sufficiently balanced in the apical dendrites that the Ca^2+^ pump was able to reduce the concentration. Somatic Ca^2+^ again increased with brief reversal of Na^+^-Ca^2+^ exchange, but was primarily driven by influx from the dendrites, trailing the basal concentrations. With the cytosolic Ca^2+^ elevated, the ER concentrations in all regions recovered, then exceeded baseline as they moved towards equilibrium with the elevated cytosolic concentrations (Figure 5e solid line).

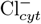 likewise exhibited a gradual increase prior to depolarization and a significant rise thereafter, primarily driven by alterations in KCC2 and NKCC1 currents, revealing differential regulation across somatic, basal, and apical regions. Prior to depolarization,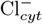 gradually increased, due to reductions in KCC2 current (Figure 5f inset). The largest increase in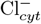 followed depolarization, driven primarily by the rise reversal of KCC2. (Figure 5f solid line). Apical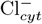 showed larger and more rapid rises due to the distribution of [K^+^]_i_ and [Na^+^]_i_. The shifts in E_K_ and E_Na_ were more abrupt near the soma due to the greater reductions in O_2_, ATP production, and Na^+^-K^+^ pump activity.

Under normoxic and baseline [K^+^]_o_ (control) conditions,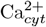 or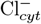 remained stable (Figure 5d&f dashed line). However, there was a gradual decline in ER Ca^2+^ concentration (Figure 5e dashed line).

### 3.4 Basal dendrites show greatest vulnerability to concentration changes

To understand the particular vulnerability of dendrites (Hartings et al., 2017), we further explored differences in voltages and fluxes from soma to both apical and basilar dendrites. Here we compared hypoxic and normoxic conditions with baseline [K^+^]_o_ (Figure 6). The activity of Na^+^-K^+^ pump is crucial for maintaining ion homeostasis; its failure under ischemic conditions gives rise to specific dendritic vulnerabilities due to neuronal morphology. Na^+^-K^+^ pump activity was driven by excess [Na^+^]_i_, [K^+^]_o_ and available ATP. Inward somatic Na^+^ currents per unit area were far larger than dendritic currents, but the resulting substantial and rapid increase in [Na^+^]_i_ was similar in the dendrites due to the greater surface/volume ratio (Figure 6a&(b). While the increase in [Na^+^]_i_ was similar throughout the neuron, the rise in [K^+^]_o_ was more prominent around the soma and basal dendrites (Figure 6c). The extracellular K^+^ did not saturate the Na^+^-K^+^ pump and so determined the maximum drive, reaching 50% of maximum first at the soma, then at the basal dendrite, while only every reaching 48% of maximum in the apical dendrite. Na^+^ influx was driven by a feedback loop, where efflux declined due to a reduction in Na^+^-K^+^ pump from the loss of ATP, and the influx increased principally via the leak currents due to shifts in the Nernst potentials.

**Figure 6.**
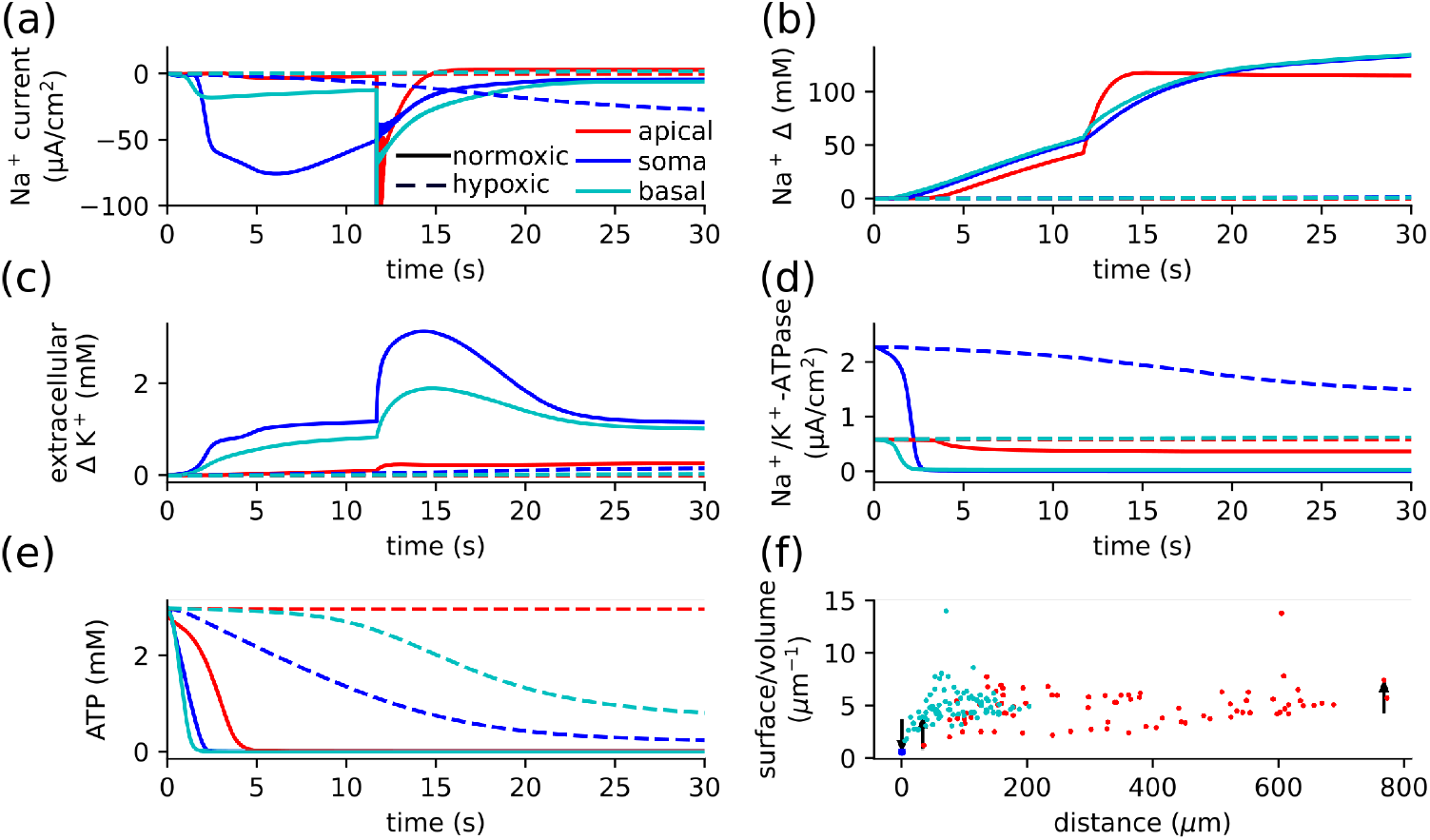
The dense dendritic arbor around the soma leads to greater vulnerability to hypoxia in the basal dendrites. **(a)** In normoxic conditions, there is little change in Na^+^ currents. Under hypoxic conditions, however, there were substantial sustained inward Na^+^ currents. **(b)** Hypoxia caused a rise in [Na^+^]_i_ from baseline throughout the neuron. Prior to depolarization (∼ 12 s) the rise is most rapid in the basal dendrites and soma, following depolarization the rise is greatest in the apical dendrites. [Na^+^]_i_ then plateaus in the apical dendrites, leading to higher concentration in the soma and basal dendrites. **(c)** Major increase in extracellular K^+^ in the soma and basal dendrites due to the large dendritic arbor near the soma. There is only a small rise at the apical dendrites. **(d)** The Na^+^-K^+^ pump current declined in the soma (falling to 10% at 2.39 s) and basal dendrites (10% at 2.07 s) under hypoxia due to the consumption of ATP. The pump was spared in apical dendrites, remaining at 62.37% by 60 s. **(e)** ATP was more rapidly consumed in the basal dendrites due to both earlier onset of Na^+^-K^+^ pump drive (due to Na^+^ influx) and larger current (due to extracellular K^+^), so they depolarized before the rest of the neuron. While ATP was consumed in the normoxic condition, the difference did not substantially impact the action of the Na^+^-K^+^ pump. **(f)** The surface/volume ratio in the apical and basal dendrites are similar, but the distance from the soma and the dense basal dendritic arbor play a role in determining the rise in extracellular K^+^. Arrows indicate the parts of the neuron compared above.

The Na^+^-K^+^ pumpactivity was determined both by demand ([Na^+^]_i_, [K^+^]_o_) and the supply of O_2_, leading to regional differences in activity and ion concentrations. There was a marked decline in Na^+^-K^+^ pump activity at the soma and basal dendrites, with apical dendrites spared (Figure 6d). The Na^+^-K^+^ pump currents depended on both the drive and ATP production, which in turn depended on available O_2_. Hypoxia led to ATP loss throughout the cell, but was most pronounced in the soma then the basal dendrite while the apical were relatively spared (Figure 6e). Despite substantial ATP loss in the apical Na^+^-K^+^ pump, there was a large residual current in the apical dendrites. This Michaelis-Mention formulation of Na^+^-K^+^ pump ATP-dependence gave a 69.2 fold reduction in Na^+^-K^+^ pump current. This reduction was counterbalanced with increased demand (elevated [Na^+^]_i_ and [K^+^]_o_), which in the absence of ATP constrains would give a 108.5 fold increase in Na^+^-K^+^ pumpactivity. Currents in the neuron depend both on the mechanism and the area of the membrane, while concentrations depend on the volume of the compartment. We consider the surface/volume ratio of each compartment Figure 6f. The compartment sizes were similar for basal and apical dendrites, with a difference driven by the density of compartments. The greater and more rapid rise in extracellular K^+^ and fall in O_2_ in the basal dendrite was due to the dense dendritic arbor and close proximity to the soma. This gave rise to the larger Na^+^-K^+^ pump currents in the apical dendrites.

### 3.5 Oxygen dependence of ischemic depolarization is all-or-none; concentration changes are gradual

With baseline K^+^ concentrations, there was an abrupt transition in membrane potential between oxygen levels that could and could not maintain the RMP, consistent with an all-or-none SD process (Figure 7a dashed-lines). Depolarization occurred earlier at lower oxygen levels, suggesting faster SD propagation with more severe oxygen deprivation. Depolarization was abrupt, but intermediate states persisted without such large pathological changes in Ca^2+^ and Cl^−^ concentrations. The electrical stimulus produced a relatively abrupt transition between oxygen levels that could maintain the RMP (0.03 mM or more) and those that could not (0.02 mM or less). Pathological depolarization occurred between 9.5 s for 1 μM O_2_ and 30.0 s for 0.02 mM O_2_. Similar abrupt transitions occur with ATP, K^+^, and Na^+^ concentrations (Figure 7b-d dashed-lines). The residual ATP was greatest in the apical dendrites (Figure 7b, this was reflected in higher [K^+^]_i_ and lower [Na^+^]_i_ due to residual Na^+^-K^+^ pump activity (Figure 7c-d). This in turn led to the greater accumulation in Ca^2+^ in the basal dendrites (primarily due to [Na^+^]_i_ depended Na^+^-Ca^2+^ exchange) and greater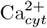 in the apical dendrites (due to E_K_ depended KCC2). With 0.03 mM O_2_ (while the neuron maintained RMP), there was a substantial reduction in ATP and subsequent partial loss of [K^+^]_i_ and accumulation of [Na^+^]_i_. These intermediate states persisted and led to a less dramatic increase in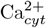 and did not produce secondary pathological changes in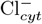 (Figure 7e&f).

**Figure 7.**
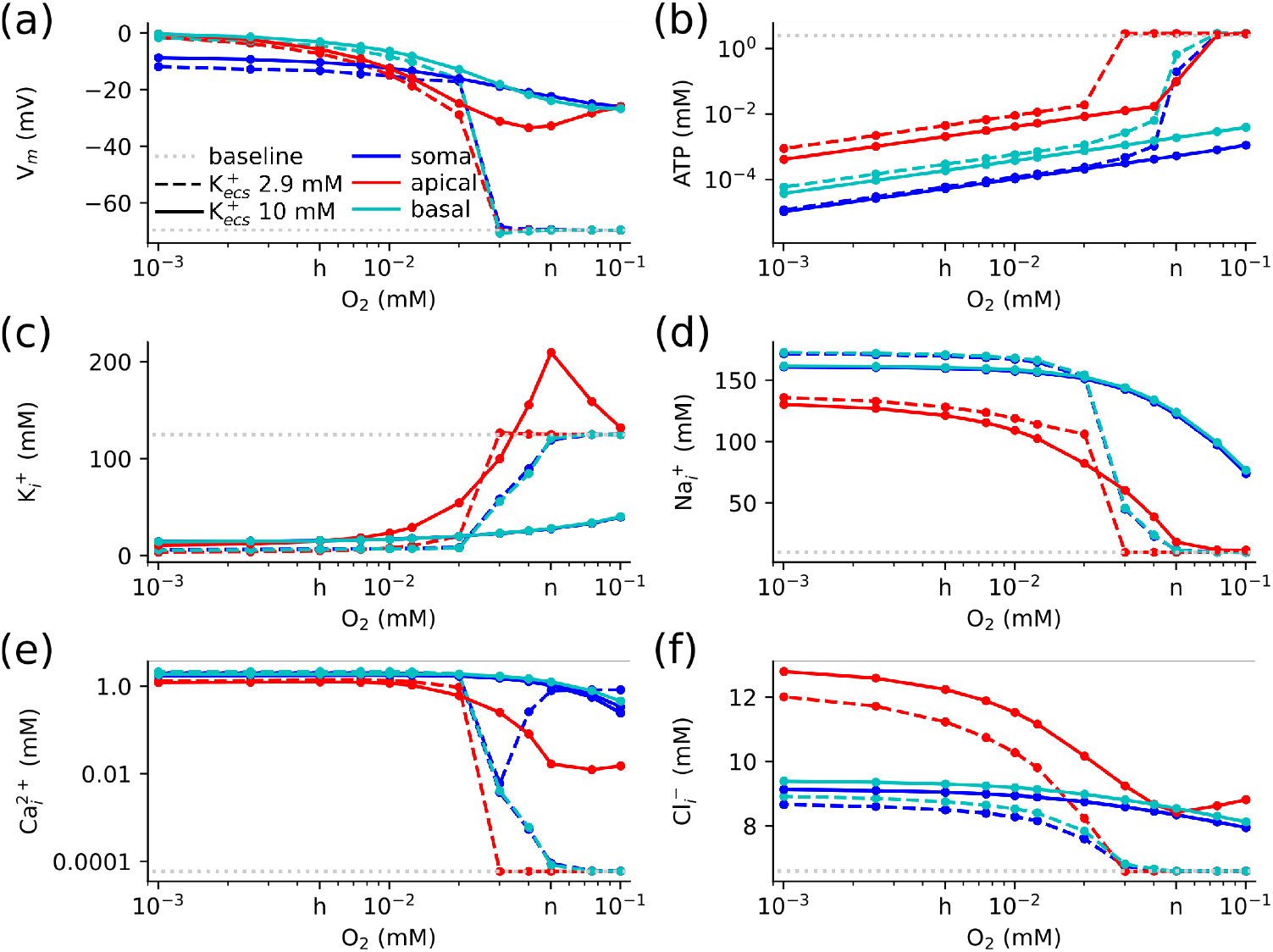
All-or-none depolarizations occurred when O_2_ was limited or extracellular K^+^ was elevated. Concentration changes depended on the extent and nature of the stressor. With elevated extracellular K^+^ (10 mM) depolarization occurred rapidly (within 30 ms) at all O_2_ levels considered (10^−3^ to 0.1 mM). In the absence of elevated K^+^ (2.9 mM) at and just below normoxic (‘n’ on the *x*-axis) O_2_ levels, the neuron maintained RMP; at lower or hypoxic (‘h’) levels, depolarization occurred after a delay (up to 30 s). This figure shows the relation between the limited O_2_ supply and: **(a)** Membrane potential, **(b)** ATP, **(c)** K^+^, **(d)** Na^+^, **(e)**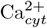, **(f)**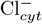.

While the dynamics were different, the pathological response to hypoxia tended to a similar state for both elevated and baseline K^+^, with substantial difference only at higher O_2_ levels (Figure 7). Elevated K^+^ produced rapid depolarizations under all O_2_ levels (Figure 7a solid-lines). With sufficient O_2_ (above 0.03 mM), apical [K^+^]_i_ increased above baseline due to large sustained Na^+^-K^+^ pump currents. Accumulation peaked at 0.005 mM O_2_, due to the effects of Na^+^ channel activation leading to higher [Na^+^]_i_ and greater Na^+^-K^+^ pump drive (Figure 7d solid-lines). Depolarization caused by extracellular K^+^ led to Ca^2+^ accumulation, which primarily depends on the glutamate mediated NMDAR influx and Na^+^-Ca^2+^ exchange clearance, which was greatest in the basal dendrites (Figure 7e solid-lines). The accumulation of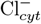 was a direct result of the shift in E_K_ via KCC2 reversal, which was greatest in the apical dendrites (Figure 7f solid-lines).

## 4 Discussion

Our simulation of a neuron under hypoxic vs. normoxic conditions gave insights into the mechanisms underlying cellular and subcellular responses to energy deficit or ionic overload, with implications for the effects at the level of bulk tissue and of the neuronal network.

The model predicted three stages of response to hypoxic loss of homeostasis: (1) replacement of [K^+^]_i_ with [Na^+^]_i_ with little initial effect on membrane potential; (2) rapid depolarization of neurons due to opening of the Na^+^ channels, particularly in dendrites with the greatest ion concentration change; and (3) persistent ion concentration breakdown. By comparison, elevated [K^+^]_o_without hypoxia led to (1) rapid depolarization due to the change E_K_; (2) the shift in Nernst potential dwarfed the synaptic mechanisms; (3) leading to different distributions of V_m_ and ions in the dendrites. The different consequences depending on the specifics of the relative severity of ionic and hypoxic insult suggest different neuroprotective therapies based on the pattern of pathophysiological patterns.

Our model demonstrated the influence of neuronal morphology on dendritic stressors due to regional differences in extracellular accumulations and local O_2_ consumption. Specifically, the density of the basal dendritic arbor near the soma is a region of high vulnerability due to (1) reduced clearance of glutamate with activation of NMDAR leading to Ca^2+^ influx; (2) greater [K^+^]_i_ and [Na^+^]_i_ due to higher [K^+^]_o_; (3) greater local competition for O_2_. We predict that high Ca^2+^ in basilar dendrites would produce excitotoxicity with earlier loss of these dendrites. By contrast, hypoxic disruption of E_K_ and E_Na_ resulted in greater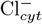 in the apical dendrites due to KCC2 imbalance. We predict that this would produce greater dendritic beading in apical as compared to basilar dendrites (Steffensen et al., 2015). This asymmetric localization of damage will have differential cognitive consequences due to the importance of apical dendrites in higher-level feedback with predominance of basilar dendrites in local activity spread.

Our study has added a focus on extracellular transmission, ion homeostasis, and a more detailed model of calcium compartmentalization and dynamics compared to prior studies. Prior modeling work utilized simplified morphologies and focused on conditions for SD initiation, neuron-glia interactions, and their contribution to seizure-like activity Kager et al. (2002, 2007); Somjen et al. (2009). These models featured electroneutrality and volume dynamics, maintained by balancing changing anion concentrations with shifts in Cl^−^and changes in osmolarity, using extracellular K^+^ alone in SD initiation, with greater astrocyte buffering delaying or preventing SD. Unlike the model presented here, these studies only considered lateral diffusion in a thin layer surrounding the cell membrane, so there was no influence from the dense dendrite arbor near the soma. The model presented here does not include electroneutrality or volume dynamics; instead, Cl^−^ was modeled independently with specific mechanisms responsible for its regulation.

Experimental observations on the speed of dysregulation following oxygen glucose deprivation are on the order of minutes to hours (Martínez-Sánchez et al., 2004; Calabresi et al., 1999) The more rapid breakdown shown in our model is likely due to the simplifications inherent in our metabolic model. Biologically, multiple mechanisms are employed to maintain ATP levels under oxygen deprivation, including glycolysis, lactate production, and mitochondrial adaptations. Anaerobic glycolysis, utilizing lactate provided by glia via the neuron lactate shuttle, offers an additional supply of ATP despite reduced O_2_ (Yellen, 2018). This lactate shuttle may be important for the post-ischemic survival of neurons (Boumezbeur et al., 2010). While we did not explicitly consider any of these mechanisms as additional sources of ATP, they would serve to shift the O_2_ dependency curve to right (Figure 7).

Each cell is embedded in tissue, the neuropil, so that extracellular fluid would be strongly influenced by other cells nearby, which will be experiencing SD as well. This could change the observed differential effects between apical and basal dendrites, depending on the degree of alignment of dendritic elements, far more aligned in hippocampus than in areas of neocortex. Hypoxic simulations with elevated [K^+^]_o_, both with and without glutamate release, also produced the same distance-dependent relationships, with peak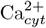 in the basal dendrites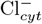 in distal-apical dendrites. The persistence of these patterns in a model of SD suggests feedback, and the cell morphology may still produce these ion gradients even when the extracellular concentrations are perturbed by its neighbors.

We note, however, that ion concentration changes during ischemia are not driven solely by a breakdown in the homeostatic mechanisms modeled here; for example, in ischemia, a rapid influx of CSF brings additional Na^+^, contributing to cytotoxic edema (Mestre et al., 2020).

Intracellular Ca^2+^ accumulation was driven primarily by NMDAR-mediated influx. When glutamate was removed from simulations, so there was no NMDAR activity,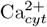 was still elevated due to the effects of Na^+^-Ca^2+^ exchange. Experiments have shown that extracellular Ca^2+^ falls to 10% of control during ischemia, with most of the uptake attributed to neurons, with estimated rises in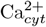 on the order of 100 nM (Silver and Erecińska, 1990; Kristián and Siesjö, 1998). Our model predicted excessive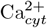 changes relative to experimental observations; this excess was due to modeling a single cell; in reality, the 0.028 mm^3^ space modeled would be occupied by ∼2,500 cells, sharing available extracellular Ca^2+^ between them.

Reducing the available extracellular Ca^2+^ in the model produced limited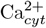 increases, consistent with experimental estimates. These low-Ca^2+^ simulations did not affect the distance-dependent patterns or the mechanisms that gave rise to them.

Varicosities distributed along neurites (‘beading’) have been observed in a variety of neuropathologic conditions, including trauma (Harris et al., 2023), epilepsy (Swann et al., 2000), neurodegeneration, and ischemia (Hori and Carpenter, 1994; Murphy et al., 2008). Specifically, in neurons in the penumbra of an experimentally induced infarct in a mouse, dendritic beading morphology was temporally correlated with SD, with recovery between episodes until a terminal injury resulting in irreversible morphologic change (Risher et al., 2010). Our results provide a mechanism by which SD links the osmotic imbalances to the beaded morphology in observed in ischemic lesions.

Additional unmodeled aspects of cellular morphology are anticipated to be consistent with the direction of changes observed in the model. Mitochondria occupy a greater fraction of intracellular volume in the basal and oblique dendrites than in the rest of the cell (Lee et al., 2022), but as the limiting factor of ATP production during ischemic is O_2_, the distribution of mitochondria would not negate the dendritic vulnerability predicted in this model. Cellular swelling due to osmotic pressure would likely dilute intracellular concentrations while reducing extracellular volume, leading to increased extracellular concentrations.

The model described here offers, to our knowledge, the most biophysically detailed model to date of the combined effects of ionic imbalance and ischemia at the subcellular level. This model will be used in future studies by coupling this in network simulations. This offers the potential for new insights into cellular response to a variety of insults – primary ischemic, migraine, inflammatory, traumatic and neoplastic with new opportunities for exploring the effects of neuroprotective intervention to prevent damage beyond the primary site of injury.

## Acknowledgments

Research supported by NIH grant R01MH086638. We thank the Yale Center for Research Computing for guidance and use of the research computing infrastructure, specifically the McCleary cluster.

## Author contributions

A.J.H.N developed the model under the supervision of W.W.L and R.A.M. M.D performed the neuropathologic evaluation. All authors contributed to the text of the manuscript.

## Supplementary information

The source code for this model is available at ModelDB modeldb.science/2017004 (access code: review).

